# A silencer repressing redundant enhancer activities revealed by deleting endogenous *cis*-regulatory element of *ebony* in *Drosophila melanogaster*

**DOI:** 10.1101/2021.03.25.436947

**Authors:** Noriyoshi Akiyama, Shoma Sato, Kentaro M. Tanaka, Takaomi Sakai, Aya Takahashi

## Abstract

The spatiotemporal regulation of gene expression is essential to ensure robust phenotypic outcomes. Pigmentation patterns in *Drosophila* are formed by the deposition of different pigments synthesized in the developing epidermis and the role of *cis*-regulatory elements (CREs) of melanin biosynthesis pathway-related genes is well-characterized. These CREs typically exhibit modular arrangement in the regulatory region of the gene with each enhancer regulating a specific spatiotemporal expression of the gene. However, recent studies have suggested that multiple enhancers of a number of developmental genes as well as those of *yellow* (involved in dark pigment synthesis) exhibit redundant activities. Here we report the redundant enhancer activities in the *cis*-regulatory region of another gene in the melanin biosynthesis pathway, *ebony*, in the developing epidermis of *Drosophila melanogaster*. The evidence was obtained by introducing an approximately 1 kbp deletion at the endogenous primary epidermis enhancer (priEE) by genome editing. The effect of the priEE deletion on pigmentation and on the endogenous expression pattern of a *mCherry*-tagged *ebony* allele was examined in the thoracic and abdominal segments. The expression level of *ebony* in the priEE-deleted strains was similar to that of the control strain, indicating the presence of redundant enhancer activities that drive the broad expression of *ebony* in the developing epidermis. Additionally, the priEE fragment contained a silencer that suppresses *ebony* expression in the dorsal midline of the abdominal tergites, which is necessary for the development of the subgenus *Sophophora*-specific dark pigmentation patterns along the midline. The endogenous expression pattern of *ebony* in the priEE-deleted strains and the reporter assay examining the autonomous activity of the priEE fragment indicated that the silencer is involved in repressing the activities of both proximal and distant enhancers. These results suggest that multiple silencers are dispensable in the regulatory system of a relatively stable taxonomic character. The prevalence of other redundant enhancers and silencers in the genome can be investigated using a similar approach.

**Author summary:** Genes are expressed at the right timing and place to give rise to diverse phenotypes. The spatiotemporal regulation is usually achieved through the coordinated activities of transcription-activating and transcription-repressing proteins that bind to the DNA sequences called enhancers and silencers, respectively, located near the target gene. Most studies identified the locations of enhancers by examining the ability of the sequence fragments to regulate the expression of fused reporters. Various short enhancers have been identified using this approach. This study employed an alternative approach in which the previously identified enhancer that regulates expression of *ebony* (a gene involved in body color formation) was deleted in a fruitfly, *Drosophila melanogaster*, using the genome-editing technique. The knockout of this enhancer did not affect the transcription level of the gene to a large extent. This indicated the presence of transcription-activating elements with redundant functions outside the deleted enhancer. Additionally, the transcription of *ebony* at the midline of the abdomen, which is repressed in the normal flies, were derepressed in the enhancer-deleted flies, which indicated that the deleted enhancer fragment contained a silencer that negatively regulates multiple enhancer activities in a spatially restricted manner.

## Introduction

The spatiotemporal regulation of gene expression during the development of organisms results in diverse phenotypes. The *cis*-regulatory elements (CREs) are nucleotides that differentially modulate the transcription levels of specific genes typically in an allele-specific manner. The most common CREs, enhancers and silencers, are located within a certain distance from the transcription start sites of the target gene and contain binding sites for the transcription activators or repressors [1,2]. Conventionally, the CREs comprising enhancers and silencers, which are unique to each expression unit, are considered as subsets of modular structures in the upstream region of a gene [3–7]. While this view reflects a fundamental principle since distinct CREs typically consist of clustered binding motifs, there are recent evidences supporting a more complex picture. Various enhancers are reported to exhibit functional redundancy or to cooperatively define the expression site boundaries [8–14]. Also, a number of enhancers with pleiotropic functions have been reported (reviewed in [15]).

The CREs in genes involved in body pigmentation pattern have been well-characterized and multiple modular enhancers that activate transcription in different body regions have been documented in detail (reviewed in [16,17]). For example, the distinct CREs of *yellow* that activate transcription in bristles, wing and body, and abdomen have been identified [18–22]. However, a recent study revealed that many sequence fragments in the regulatory region of *yellow* exhibit redundant and cryptic enhancer activities, suggesting that *cis*-regulatory modules are not as distinct as described previously and more amenable to evolutionary changes [23].

A complex architecture of *cis*-regulatory region has also been implicated from the within-species comparisons of *cis*-regulatory sequences of *ebony*, another gene involved in body pigmentation. Polymorphisms in *ebony*, which encodes an enzyme of the melanin biosynthesis pathway, is the major causative factor determining the body pigmentation intensity in *D. melanogaster* [24–27]. The sequence polymorphisms of the primary epidermis enhancer (priEE), which was identified to be located in the upstream intergenic region of *ebony*, were analyzed in detail. Some single-nucleotide polymorphisms (SNPs) first identified in the African populations affected the enhancer function but were not associated with body pigmentation intensity in the Japanese, European, North American, and Australian populations [27–32]. Also, a priEE haplotype associated with light body color was identified in the Iriomote and Australian populations but not in the African populations [32]. Furthermore, there were no SNPs or indels in the priEE that showed complete association with the allele-specific expression levels in the developing epidermis in 20 strains sampled from the *Drosophila melanogaster* Genetic Reference Panel originated from a natural population in North Carolina [28,33]. Moreover, some strains with identical priEE sequences exhibited different allele-specific expression levels. These analyses suggest the possibility of the presence of sequences outside the priEE region that regulate the expression level of *ebony*.

The priEE segment was the only segment within the approximately 10 kbp regulatory region (including an upstream intergenic region and intron 1) that drove the expression of the reporter gene in the epidermis [26]. However, the previous findings above have indicated that the sequence variation within the priEE was not sufficient to explain the wide range of expression level variation of this gene. Therefore, we hypothesized that similar to *yellow, ebony* has redundant enhancers in addition to the priEE, possibly located outside the known approximately 10 kbp regulatory region, and may be taking part redundantly in activating transcription of this gene in the developing epidermis.

The expression pattern of *ebony* is repressed in certain areas of the abdomen of *D. melanogaster*, which results in the generation of a distinct pigmentation pattern [26,34]. As the transcription of *ebony* is simultaneously activated by multiple enhancers, silencing mechanisms must be present to suppress the simultaneous activation. In *D. melanogaster*, dark stripes are visible in the posterior regions of the A2–7 abdominal tergites in females and the A2–4 tergites in males. Also, the abdominal pigmentation pattern exhibits sexual dimorphism with totally dark A5–7 tergites observed only in males. Furthermore, another characteristic dark line is present along the dorsal midline of the abdominal tergites in both males and females of this species. The dark line is a characteristic of the subgenus *Sophophora* [35] and the expression of *ebony* is repressed in this region [26,34]. The locations of silencers responsible for stripe repression and male-specific repression in the posterior tergites have been suggested [26]. However, a silencer for repression at the dorsal midline has not been identified previously.

The transgenic reporter assay is a powerful approach to dissect regulatory sequences and identify CREs, such as enhancers and silencers. However, some limitations exist because this assay does not test the sequence fragments in their native genomic environment [36]. Especially, the lengths and borders of the sequence fragments can markedly affect the results [23]. In this study, rather than conducting reporter gene assays, we examined the genomic region by modifying the endogenous upstream sequence using the clustered regularly interspaced short palindromic repeats (CRISPR)-CRISPR associated protein 9 (Cas9) system. The precise knockout of the known priEE in a clean genetic background enabled the examination of its contribution to the body color phenotype. We combined it with an assay using a reporter gene tagged to the endogenous *ebony* to capture the changes in the expression pattern. As a result, we uncovered the presence of redundant enhancer activities that drive the broad expression of *ebony* in the developing epidermis. We also show that the priEE fragment contains silencers for repressing the expression of *ebony* in the dorsal midline of the abdominal tergites, which is necessary for developing the *Sophophora*-specific pigmentation pattern. This silencer represses the activities of the proximal and distant enhancers. We discuss the consequences of such regulatory system on the evolution of CREs and the potential application of a similar approach to other genomic regions.

## Results

The priEE fragment was precisely knocked out using the CRISPR-Cas9 system to examine whether transcriptional activation of *ebony* occurs in the absence of the priEE. First, to control the genomic background, an isogenic Cas-0002 line (Cas-0002-iso) carrying the *nos*-Cas9 transgene was constructed (Figs 1A and S1). Next, *mCherry* was knocked-in to the 3’ end of *ebony* in the Cas-0002-iso line using the CRISPR-Cas9 system (Fig 1B). The resultant transgenic line (Cas-0002-iso_*e*::*mCherry*) was designed to produce a Ebony-mCherry fusion protein. Cas-0002-iso and Cas-0002-iso_*e*::*mCherry* lines were crossed with guide RNA (gRNA) expression lines to drive a targeted deletion at the approximately 970-bp priEE fragment [28,29] (Fig S2). Additional crosses were performed to remove *y*^*2*^ and replace the X chromosome with *w*^*1118*^ to avoid interference from *yellow*, which is in the same pigment synthesis pathway (Fig S2). The following three priEE deletions were generated; two from Cas-0002-iso line (*w*^*1118*^; *e*^*Δ1088priEE*^ *and w*^*1118*^; *e*^*Δ1089priEE*^) and one from Cas-0002-iso_*e*::*mCherry* line (*w*^*1118*^; *e*^*Δ1077priEE*^::*mCherry*) (Figs 1C–D). If the priEE contains the only enhancer driving the expression of *ebony* in the epidermis, the priEE-deleted strains must exhibit a dark body color equivalent to the *ebony* null mutant (*e*^*1*^). Contrary to this prediction, the body color of the priEE-deleted strains was similar to that of the control strain (Fig 2A).

**Fig 1.**
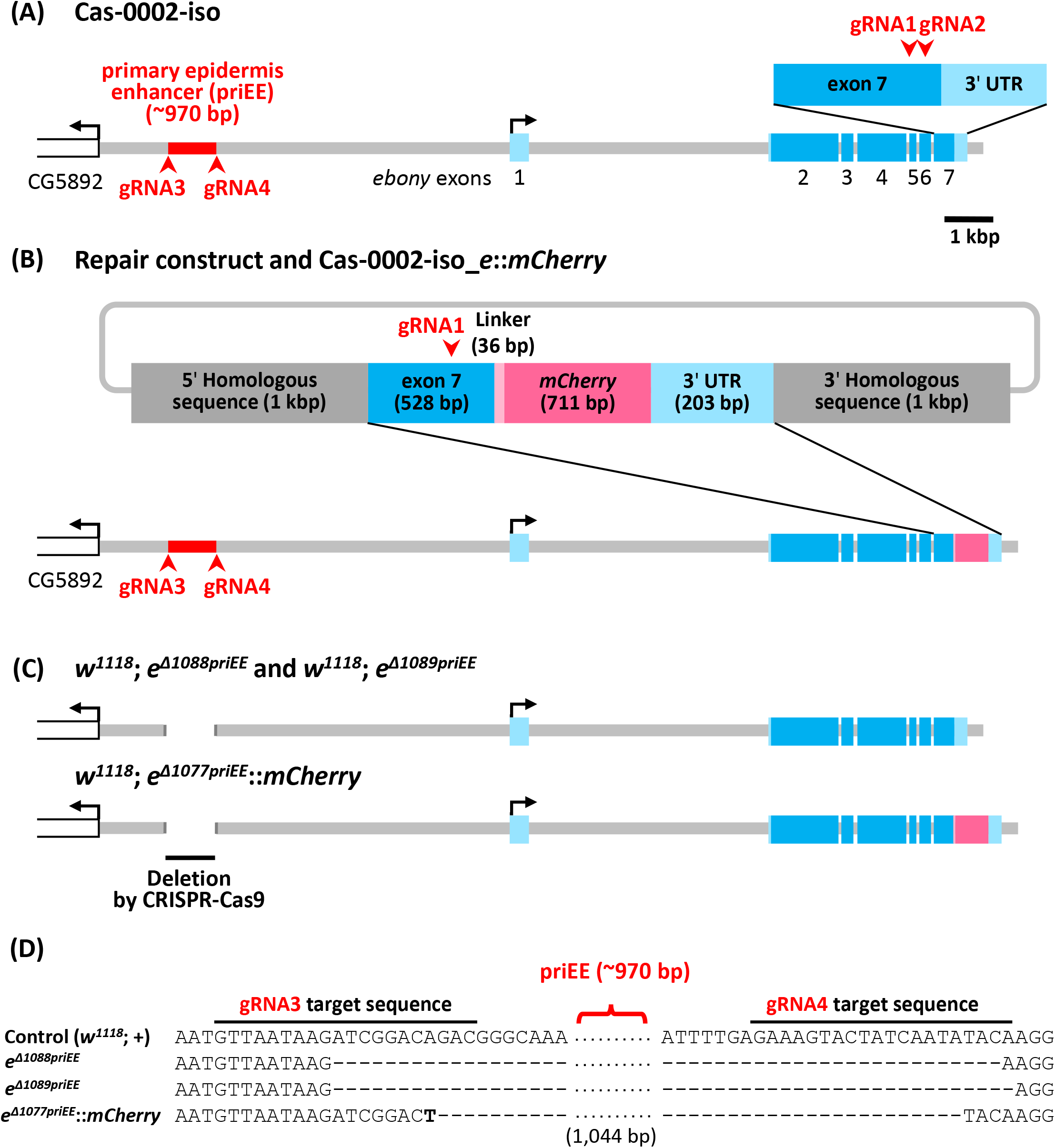
Construction of the primary enhancer element (priEE) knockout strains. (A) The genomic region surrounding *ebony* in Cas-0002-iso, an isogenic line carrying *nos*-Cas9. (B) The genomic region surrounding *ebony* in Cas-0002-iso_*e*::*mCherry* is shown with the repair construct for *mCherry* knock-in. (C) The genomic region surrounding *ebony* in strains with deleted priEE after the removal of *y*^*2*^ (Fig S2). The light blue box indicates the untranlated region (UTR) and the blue box indicates the coding sequence (CDS). The red arrowhead indicates the target site of guide RNA (gRNA) sequences. (D) Partial sequence alignment around the priEE fragment in the control and priEE-deleted strains. *w*^*1118*^; *e*^*Δ1077priEE*^::*mCherry* had a single T of unknown origin within the deleted region.

**Fig 2.**
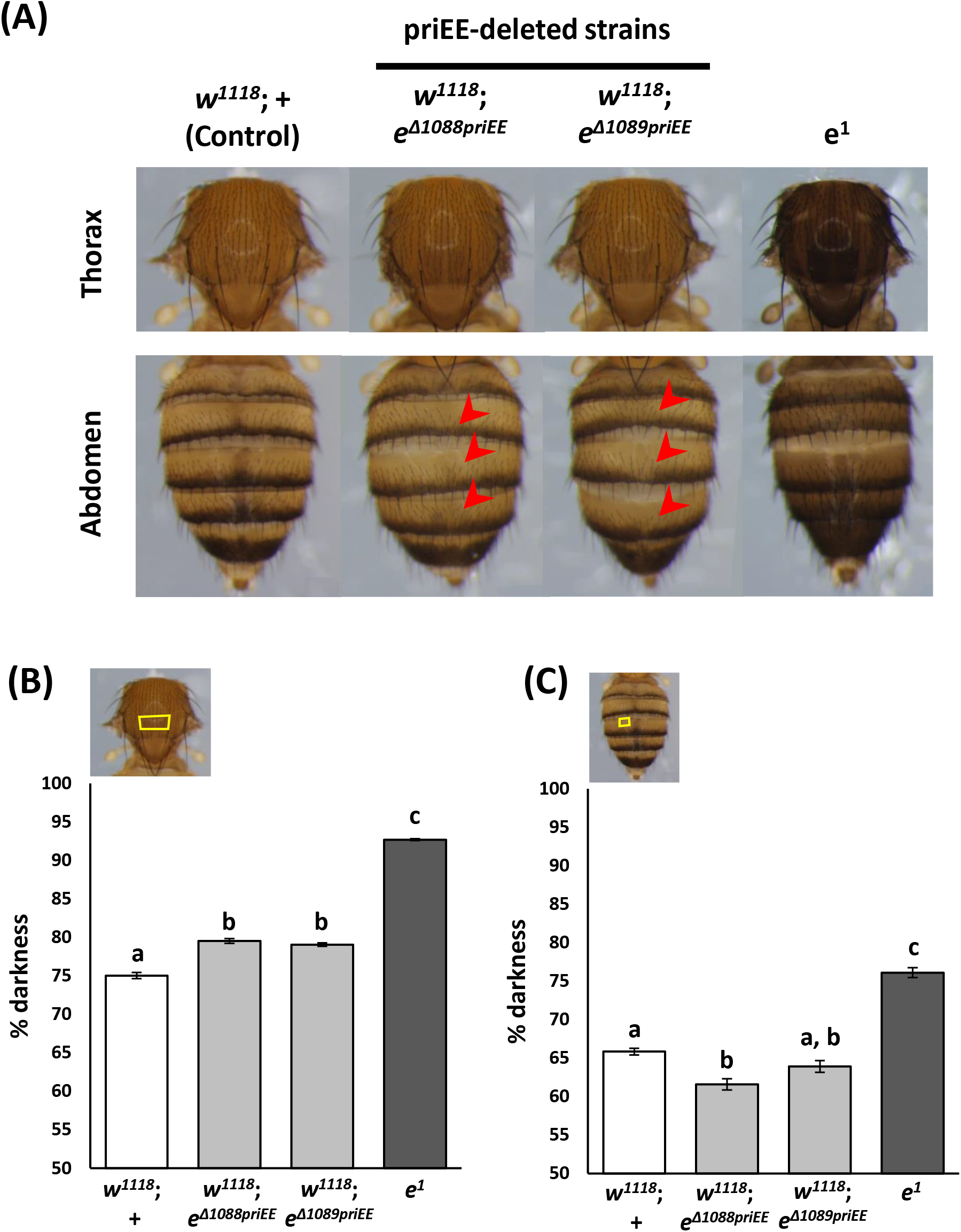
Effect of primary enhancer element (priEE) knockout on the intensity and patterns of pigmentation. (A) Images of the 5–7-day-old adult females. The red arrow indicates the area where the dark pigmentation in the dorsal midline is missing. (B) Percent (%) darkness values of the thorax measured at the area enclosed in the yellow square. (C) Percent (%) darkness values of the A4 abdominal segment measured at the area enclosed in the yellow square. *N* = 10 for each strain. Different letters indicate significant differences between strains (*P* < 0.05; one-way analysis of variance, followed by Tukey HSD post-hoc test). Error bars denote standard error.

The pigmentation intensity in the females of the two priEE-deleted strains (*w*^*1118*^; *e*^*Δ1088priEE*^ *and w*^*1118*^; *e*^*Δ1089priEE*^), the control strain (*w*^*1118*^; +), and an *ebony* null mutant (*e*^*1*^) was compared. The percent darkness at the specific positions of the thoracic center and the fourth abdominal tergite (A4) was measured (10 flies per strain) (Fig 2B–C). The pigmentation scores were significantly different between the strains (thorax, *F*_3, 36_ = 733.59, *P* < 10^−15^, one-way analysis of variance (ANOVA); abdomen, *F*_3, 36_ = 94.919, *P* < 10^−15^, one-way ANOVA). The thoraces of the two priEE-deleted strains showed significantly but only slightly darker pigmentation than those of the control strain (Fig 2B). The abdominal pigmentation in the *w*^*1118*^; *e*^*Δ1088priEE*^ strain was slightly lighter than that in the control strain, but that in the *w*^*1118*^; *e*^*Δ1089priEE*^ strain was not significantly different from the control strain (Fig 2C). These subtle changes in pigmentation intensity suggest that the priEE deletion perturbs *ebony* transcription to some extent. However, the pigmentation in the thorax and abdomen of the two priEE-deleted strains was markedly lighter than that in the thorax and abdomen of *e*^*1*^ (Figs 2B–C). This indicated that deletion had limited effects on the overall transcription level regulation. Hence, the priEE was shown to be dispensable for driving the transcription of *ebony* in the developing epidermis.

Also, unexpectedly, the deletion of the priEE affected the pigmentation pattern. In particular, the deletion of the priEE resulted in the loss of a dark pigmentation line along the dorsal midline of the abdominal tergites (Fig 2A). This indicated that *ebony* expression is suppressed in the midline area and that the priEE fragment is necessary for this suppression.

To directly investigate the expression sites of this gene in the abdomen, the abdominal epidermis of the *mCherry* knocked-in strains, *w*^*1118*^; *e*::*mCherry* and *w*^*1118*^; *e*^*Δ1077priEE*^::*mCherry* (Fig 1C), were subjected to fluorescence confocal microscopy (Figs 3A–B). The abdominal pigmentation of *w*^*1118*^; *e*^*Δ1077priEE*^::*mCherry* was consistent with the priEE-deleted strains without *mCherry* (*w*^*1118*^; *e*^*Δ1088priEE*^ *and w*^*1118*^; *e*^*Δ1089priEE*^) (data not shown), which suggested that the catalytic function of Ebony in the pigmentation synthesis pathway is not disrupted upon fusion with mCherry. Endogenous *ebony* exhibited a broad epidermal expression pattern (as expected from the light pigmentation), and a suppressed expression at the posterior stripe region of each tergite (A1–6 of a female and A1–4 of a male) and tergite-wide suppression at A5 and A6 in males (Figs 3A, D, and E). These expression patterns were consistent with those reported in previous studies [17,34]. As predicted from the pigmentation scores in Fig 2A, *ebony* expression in *w*^*1118*^; *e*^*Δ1077priEE*^::*mCherry* was similar to that in the control strain except that the expression at the dorsal midline was not suppressed (Figs 3B, D, and E). These results indicate that the enhancer activity resides outside the primary enhancer and that redundant enhancer element(s) is present in the surrounding genomic region. The expression of *ebony* has been reported in other body regions [26,37–39] but the priEE has not been indicated to drive expression in tissues other than the developing epidermis [26]. As expected, the expression patterns of *ebony* did not markedly change in other tissues upon deletion of the priEE (Fig S3).

**Fig 3.**
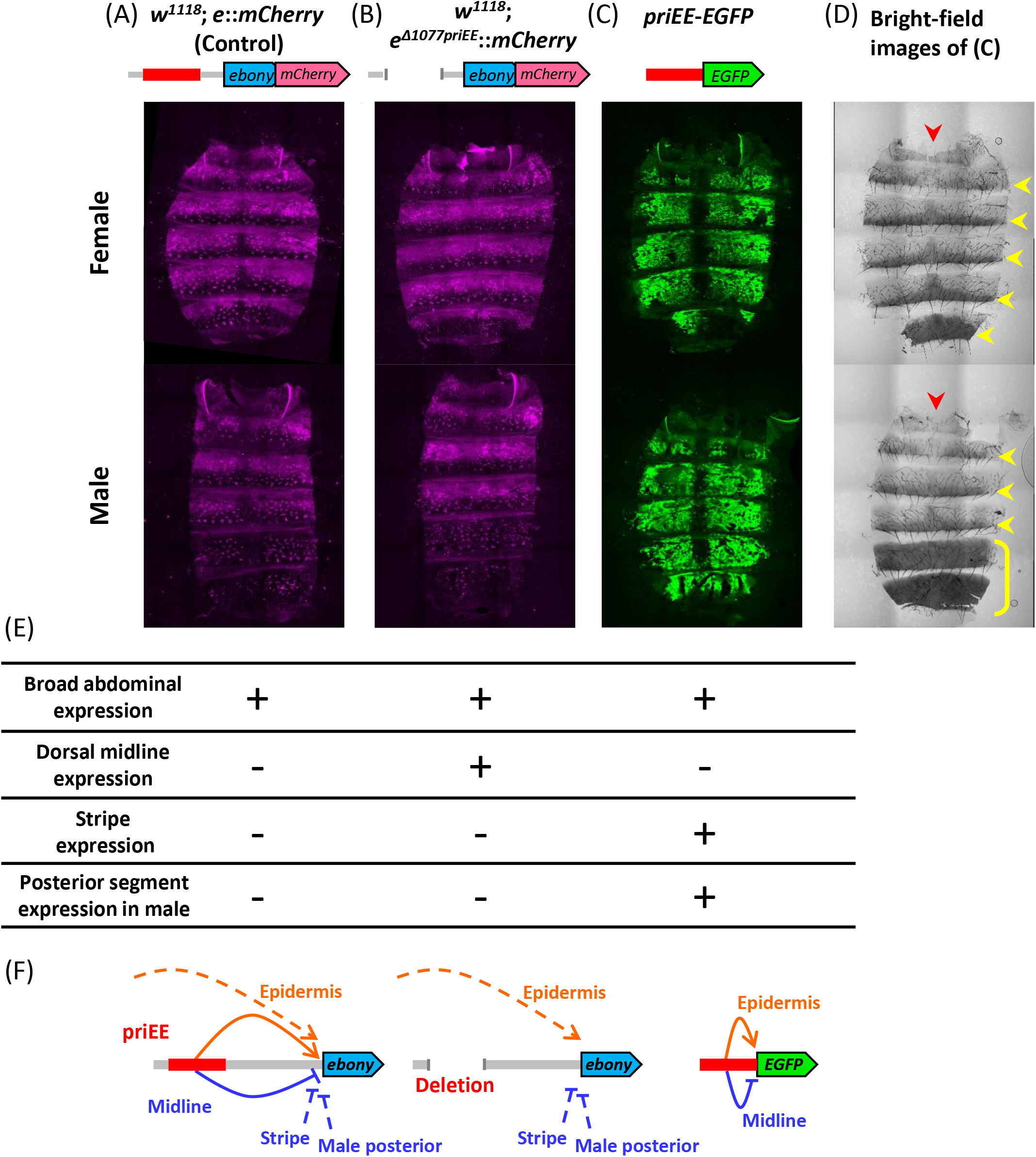
*cis*-regulatory elements (CREs) regulate *ebony* expression in the developing epidermis. Confocal fluorescence images of the developing abdominal epidermis of *w*^*1118*^; *e*::*mCherry* (A), *w*^*1118*^; *e*^*Δ1077priEE*^::*mCherry* (B), and *priEE*-*EGFP* transformed to VK00033 (C). (D) Bright-field images of (C). Images of females and males are shown in the upper and lower panels, respectively. The red arrowhead indicates the dorsal midline. The yellow arrowhead indicates a dark stripe on the posterior area of each tergite. The yellow bracket indicates male-specific dark pigmentation in the A5 and A6 tergites. (E) Summary of the *ebony* expression sites for each strain determined from the fluorescence signals. (F) The suggested model for the regulation of *ebony* expression in the abdomen. The solid lines indicate the effects of priEE, while the dotted lines indicate the effects of other redundant CREs. The priEE fragment was also equipped to function as a midline silencer.

The results indicated that the deleted priEE fragment contained a silencer element for the dorsal midline as well as an epidermal enhancer element. In order to confirm that the deleted priEE contains both elements, a GFP reporter assay was performed. The priEE fragment was fused to an enhanced GFP (*EGFP*) gene (*priEE*-*EGFP*) with a minimal Hsp70 promoter and transformed into two attP strains (VK00033 and VK00037). The confocal images of GFP from *priEE*-*EGFP* transformed to a third chromosome landing site in VK00033 (Figs 3C and E) and a second chromosome landing site in VK00037 (Fig S4) indicated that the priEE autonomously drives the epidermal expression except at the flanking regions of the dorsal midline. Rebeiz *et al*. [26] reported that the 0.7 kbp core element (included in the approximately 970-bp priEE) drove a similar expression pattern. The pattern clearly showed that a dorsal midline silencer is present in the priEE fragment and that it can silence the activity of the proximal enhancer element within the priEE fragment, which drives the broad expression of *ebony* in the developing abdominal epidermis.

## Discussion

### Redundant enhancer activity resides in the regulatory region of *ebony*

The removal of the endogenous priEE of *ebony* using the CRISPR-Cas9 system did not cause a drastic darkening as observed in the null mutant (*e*^1^), although a slight perturbation of the pigmentation intensities in the thoracic and abdominal segments was observed (Fig 2). A strong negative correlation between the darkness of body pigmentation and the expression level of *ebony* in the developing epidermis has been repeatedly detected in strains sampled from the natural populations of *D. melanogaster* [24–29]. Thus, the dark pigmentation intensity of the cuticle is suggested to be a sensitive indicator of local changes in the expression level of *ebony*. Therefore, the lack of a large increase in dark pigmentation in strains with the priEE deletion indicated that transcription of the gene was not largely disrupted. Further, these findings indicated that the transcriptional activation of *ebony* in the developing epidermis can also be driven by the endogenous sequences outside the priEE, suggesting the presence of redundant enhancer elements in the surrounding genomic region. The complex arrangement of multiple CREs may be a reason for the scarcity of polymorphisms association with pigmentation intensity or gene expression level within or near the enhancer element across worldwide populations [32].

The locations of the redundant elements have not been determined. The results of a previous reporter assay revealed that no fragments other than those including the priEE segment were detected within the approximately 10 kbp regulatory region that contains the 5’ intergenic region and the first intron [26]. Therefore, redundant enhancer elements are likely to be located elsewhere. However, unlike the recent reporter assay conducted with the *yellow* regulatory region [23], many regions were tested using relatively large fragments (> 2 kbp), which may contain cryptic enhancers that are repressed by surrounding sequences in their native genomic context. Therefore, the possibility of the presence of redundant enhancer elements within the approximately 10 kbp regulatory region cannot be ruled out. Some secondary enhancers are reported to be shadow enhancers that are more than 20 kbp away from the transcription start site [8]. Thus, there is a need for extensive search to elucidate the detailed spatial arrangement of CREs. Nevertheless, the advantage of deleting an endogenous enhancer, a strategy employed in this study, is the rapid capturing of redundant enhancer activity in the native genomic context. Such knockout assays using endogenous genome editing may reveal more cases of redundant enhancer activities in the *Drosophila* genome as in the study conducting similar experiments on mouse developmental genes [40]. Moreover, this approach can compensate for some potential bias in reporter gene assays caused by the choice of promoter and the genomic location of the transgenes [36].

### Possible functions of redundant enhancers are to be investigated

Redundant enhancer elements, which are often referred to as primary and shadow enhancers [8], have been suggested to confer robustness against environmental or genetic perturbations [10,12,39] or define sharp boundaries for gene expression [11,13]. The transcriptional activation of *ebony* by multiple enhancers appears to be largely overlapping but may not be completely redundant, considering the subtle changes in the pigmentation intensity upon deletion of the priEE (Fig 2B). However, wide range of variations in the transcription level of this gene have been reported within and among *D. melanogaster* populations [24–29]. Thus, maintaining a robust transcription level of this gene might not be essential. The functional significance of redundant enhancers in this gene requires further investigation.

### A single silencer represses the activity of multiple redundant enhancers

Spatially restricted suppression of focal gene transcription can be achieved by introducing a specific silencer that recruits repressive transcription factors (or repressors) expressed in the target cells. In contrast to enhancers, there is far limited information on the exact locations and features of silencers. This study demonstrated that the priEE fragment contained a silencer of *ebony* expression in the abdominal dorsal midline based on two experimental evidences. First, the suppression of *ebony* expression at the dorsal midline was not observed when the priEE fragment was deleted (Figs 2A and 3B). Second, experiments with priEE fragment fused to a reporter gene revealed a broad epidermal expression driven by autonomous enhancer activity and the suppression of gene expression at the dorsal midline (Fig 3C). These results also have implications about the functional category of the silencer in accordance to its effective range.

The repressors can be grouped into the following two categories: short-range repressors, locally inhibit activators at distances of less than 100–150 bp, long-range repressors, inhibit activators at distances of at least 500 bp or longer [41–43]. Generally, long-range repressors can function in a dominant fashion to block multiple enhancers [41]. The *ebony* expression in the priEE-deleted strain implies that in the wildtype strain, when the silencer is intact, multiple enhancers of this gene are simultaneously inhibited. The results of the *priEE-EGFP* reporter assay demonstrated that the repressor bound to the silencer within the fragment interferes with the proximal enhancer, and may exhibit short-range repression. Taken together, a single silencer is sufficient to inhibit all the redundant activities from the proximal and distant enhancers, possibly by interfering with the basal transcription machinery at the promoter site. Also, the repressor bound to this silencer might have a potential to directly interact with the proximal enhancer element at short-range. Various models have been described to explain the long-range and short-range functions of repressors [41–45]. A chromatin conformation analysis may be effective to identify the direct physical interaction between the silencer and the promoter.

A schematic representation of the *cis*-regulatory transcriptional control of this gene is shown in Fig 3F. As incorporated in the model, the repression of *ebony* in the dark stripes at the posterior regions of the abdominal tergites and the totally dark A5–6 tergites in males is not affected (Figs 3A–D). This is consistent with the results of a previous study, which showed that the locations of these silencers are not within the deleted fragment [26]. The authors revealed that the male silencer was located approximately 1.5 kbp upstream of the transcription start site, and the stripe silencer was located within the first intron. Whether these CREs recruit long-range and/or short-range repressors or not would be an intriguing question for obtaining a comprehensive picture of the regulatory system of this gene. A similar approach to remove the putative silencer region can be effective for the purpose.

At the molecular level, *omb, dpp*, and *wg*, are reported to be involved in the formation of sexually monomorphic pigmentation patterns in the abdomen of *D. melanogaster*, and *dpp*, which is expressed at the dorsal midline is essential for the formation of dark pigmentation along the midline [46–48]. Additionally, *dpp* is known to activate the BMP signaling pathway, which regulates the transcription of numerous genes through a downstream transcription factor Mad (reviewed in [49]). Kopp *et al*. [46] showed that *Mad*^*12*^ clones at or near the dorsal midline promoted the loss of dark pigmentation, which suggested that Dpp signaling contributes to pigmentation. Furthermore, an RNAi screening revealed that 48 transcription factors, including Mad, are involved in abdominal pigmentation [50]. Therefore, it is likely that the suppression of *ebony* by the silencer is regulated through the Dpp signaling pathway.

### Derepression of *ebony* is sufficient to diminish a taxonomic character

In the genus *Drosophila*, the pigmentation pattern of the abdominal midline is one of the traits used to classify the subgenus *Sophophora*, which includes *D. melanogaster*, and the subgenus *Drosophila*. With some exceptions, the pigmentation stripes on the abdominal tergites of the subgenus *Sophophora* are mostly connected or expanded anteriorly at the dorsal midline forming a distinct dark area along the midline as in *D. melanogaster* (Figs 2A and 3D). In contrast, the stripes are narrowed or broken at the midline in most species of the subgenus *Drosophila* [35]. We have shown that the suppression of *ebony* by the abdominal midline silencer is at least necessary for the *Sophophora*-type midline to appear in *D. melanogaster*. The expression patterns of *pale, Ddc, ebony, tan*, and *yellow* in the developing abdominal epidermis of species belonging to subgenus *Sophophora* were previously examined using *in situ* hybridization [34]. Among the investigated genes, the suppression of *ebony* appears to be most pronounced in species with a typical dark dorsal midline.

A pair of sister species within the subgenus *Drosophila, D. americana* and *D. novamexicana*, represents another case of distinct pigmentation patterns in the abdominal midline. *D. americana* has a dark body color with uniformly dark abdominal tergites, whereas *D. novamexicana* exhibits a light pigmentation along the abdominal midline [51], which is a typical pattern of the subgenus. A recent study used reciprocal hemizygosity testing to demonstrate that the difference in abdominal midline pigmentation intensity between the two species was due to *ebony* [52]. The authors showed that *ebony* is required for the development of light pigmentation along the dorsal midline in wild-type *D. novamexicana*. It has not been demonstrated whether the interspecific differences of *ebony* resides in the *cis*-regulatory region or not. However, the study also suggests that *ebony* suppression might be a key factor for determining this taxonomically important trait.

In this study, a single silencer was sufficient to suppress the activities of multiple enhancers in the *cis*-regulatory region of *ebony*. This long-range repression eliminates the need for an acquisition of repressor-binding sites for individual enhancer elements. A study of *yellow*, which is also expressed in the developing epidermis, from three different species revealed a contrasting picture of frequent evolutionary acquisition and loss of short-range repressor binding sequences [23]. Also, reporter assays examining the effect of the male-specific silencer of *ebony* in *D. auraria* and *D. serrata* observed a frequent loss of this silencer in these species [53]. The differences in silencer properties may be attributed to the evolutionary stability of the focal expression sites. The presence of a universal silencer may be prevalent in genes responsible for relatively stable taxonomic characters that delimit certain clades of species. Such silencers enable the redundant enhancer elements to fine-tune their regulation while maintaining robust transcription suppression in a spatially restricted manner.

These findings, together with the recently accumulating evidences of redundant CREs, suggest that the architectures of *cis*-regulatory regions are diverse and the possible evolutionary regimes may be more complex and variable than the conventional view of modularly restricted evolution of CREs.

## Materials and Methods

### Fly strains

*y*^*2*^, *cho*^*2*^, *v*^*1*^, P{*nos-*Cas9, *y*^*+*^, *v*^*+*^}1A/*FM7c, Kr*-GAL4, UAS-*GFP* (Cas-0002), *y*^*1*^, *v*^*1*^, P{*nos*-phiC31¥int.NLS}X; attP40 (II) (TBX-0002), *y*^*2*^, *cho*^*2*^, *v*^*1*^; *Sco*/*CyO* (TBX-0007), and *y*^*2*^, *cho*^*2*^, *v*^*1*^; *Pr, Dr*/*TM6C, Sb, Tb* (TBX-0010) lines were obtained from the NIG-FLY Stock Center. *w*^*1118*^; *wg*^*Sp-1*^*/ CyO*; *Pr*^*1*^ *Dr*^*1*^*/ TM3 Sb*^*1*^ *Ser*^*1*^ (DGRC#109551) and *e*^1^ (DGRC#106436) were obtained from the Kyoto Stock Center. The isogenized Cas-0002 strain (Cas-0002-iso) was established via the triple balancer by crossing DGRC#109551 with Cas-0002 (Fig S1). The TBX-double-balancer (*y*^*2*^, *cho*^*2*^, *v*^*1*^; *Sco*/*CyO*; *Pr, Dr*/*TM6C, Sb, Tb*) was generated from TBX-0007 and TBX-0010. The two attP strains *y*^*1*^ *w*^*1118*^; PBac{*y*^*+*^-attP-3B}VK00033 (DGRC#130419) and *y*^*1*^ *w*^*1118*^; PBac{*y*^*+*^-attP-3B}VK00037 (DGRC#130421) were used to generate transgenes at WellGenetics. All fly stocks were reared at 25°C on a standard corn-meal fly medium.

### Repair construct for *mCherry* knock-in

The repair construct (Fig 1B) was designed following the method described by Hinaux *et al*. [54]. The construct *pJet-yellow_F4mut-mCherry* was gifted from Dr. Nicolas Gompel. A part of the *ebony* locus (2,610 bp (from approximately 1 kbp upstream of exon 7, to approximately 1 kbp downstream of 3’UTR)) was amplified from Cas-0002-iso (Fig 1A) using primers with the XhoI, (5’-AGCctcgagTGGTGGATAAGGCCATTGTT-3’) and XbaI (5’-CAGtctagaTGCAACTGGTTTGTGCGTAT-3’) digestion sites. PCR was performed using KAPA HiFi HotStart ReadyMix (Kapa Biosystems). The *pJet-yellow_F4mut-mCherry* vector was digested with XhoI and XbaI and the fragments flanked by these digestion sites (including partial *yellow* and *mCherry* gene sequences) were replaced by the PCR product, which was digested with the same restriction enzymes. The complete sequence of the resulting vector, excluding the *ebony* termination codon, was PCR-amplified using the following primers: 5’-GACGACCACCCGGTGGACGT-3’ and 5’-TTTGCCCACCTCCTTCCAAT-3’.

Next, the *mCherry* sequence with a 5’ linker [55] was amplified from the *pJet-yellow_F4mut-mCherry* vector using primers with 15 bp homologous flanking sequences (5’-AAGGAGGTGGGCAAAGGATCCGCTGGCTCCGCTGCTG-3’ and 5’-CACCGGGTGGTCGTCTTACTTGTACAGCTCGTCCATGCC-3’). These two amplicons were fused using the In-Fusion HD Cloning Kit (TaKaRa) to generate the *pJet*-*ebony*-*mCherry* vector.

Finally, the two synonymous mutations were inserted at the target sequence of Grna (5’-GCGCGCTATTGTCCATTGGA-3’) to reduce the risk of the repair construct being cut during the knock-in reaction. To induce mutations, two overlapping amplicons, including the gRNA target sequence, from the *pJet-ebony-mCherry* vector were generated using PCR with the following primer pairs: 5’-AATCCCCGCGAGAACATC-3’ and 5’-TCCAGTGTACAATAGCGCGC-3’; 5’-GCGCTATTGTACACTGGAAG-3’ and 5’-TTGTCTGGAAATCAAAGGCTTA-3’. These two PCR products were connected using overlap extension PCR to generate a mutated fragment. This mutated fragment was replaced by the original homologous sequence of the *pJet-ebony_mut-mCherry* vector by fusing the mutated fragment and the PCR product amplified from the *pJet-ebony-mCherry* vector using the In-Fusion HD Cloning Kit (TaKaRa) with the following primers: 5’-GCCTTTGATTTCCAGACAA-3’ and 5’-GTTCTCGCGGGGATTCAAC-3’. The constructed *pJet-ebony_mut-mCherry* vector was used as the repair construct for *mCherry* knock-in.

### gRNA vector cloning

All the guide sequences of gRNAs were cloned into the pCFD5 vector (Addgene ##73914) according to the pCFD5 cloning protocol [56]. The guide sequences of gRNA1 (5’-GGAGCACGAGGTTCTGCGGG-3’) and gRNA2 (5’-GCGCGCTATTGTCCATTGGA-3’) were designed within exon 7 of *ebony* and cloned into separate pCFD5 vectors. The guide sequences of gRNA3 (5’-GTTAATAAGATCGGACAGAC-3’) and gRNA4 (5’-GAAAGTACTATCAATATACA-3’), which were designed at both ends of the approximately 970-bp priEE fragment (Fig S5, [28,29]), were cloned into a single plasmid. An In-Fusion HD Cloning Kit (TaKaRa) was used for cloning.

### Construct for reporter gene assay

The sequence of the priEE of *ebony* was amplified from Cas-0002-iso (Fig 1B) using the following primers with restriction enzyme digestion sites: 5’-CGGgaattcGGGCAAAGCAGGGTGAATA-3’ (EcoRI site) and 5’-ACTgcggccgcTCACAGGGACTTATGGGAAA-3’ (NotI site). These primers were designed to amplify most of the priEE knocked out sequences including the whole *e*_ECR0.9 [29] and *e*_core_*cis* [28] elements (Fig S5). The amplified product and the pEGFP-attB vector with a minimal Hsp70 promoter (*Drosophila* Genomics Resource Center) were digested with EcoRI and NotI. The PCR product was cloned into the multi-cloning site of the vector.

### Embryonic microinjection

For *mCherry* knock-in, an *ebony* knockout strain was generated by injecting the gRNA1 guide-sequence-cloned pCFD5 vector (200 ng/μl) into the embryos of Cas-0002-iso. The embryos of *ebony* knockout strain were injected with a mixture of the gRNA2 guide-sequence-cloned pCFD5 vector (200 ng/μl) and the repair construct *pJet-ebony_mut-mCherry* (400 ng/μl). Of the 250 adult flies that emerged from the injected embryos, four restored wild-type body color. The sequences of exon 7 of *ebony* and knocked-in *mCherry* were confirmed using Sanger sequencing with a BrilliantDye Terminator cycle sequencing kit (NimaGen) and an ABI PRISM 3130xl Genetic Analyzer (Applied Biosystems). The established strain was named Cas-0002-iso_*e*::*mCherry* (Fig 1B).

The pCFD5 vector with guide sequences of gRNA3 and gRNA4 (200 ng/μl) was injected into the embryos of TBX-0002. The gRNA expression strain (*y*^*2*^, *cho*^*2*^, *v*^*1*^; attP40{gRNA, *v*^*+*^}; *Pr, Dr*/*TM6c, Sb, Tb*) was established by mating the successfully transformed individual with the TBX-double-balancer. The guide sequences of gRNAs of the established strains were confirmed using Sanger sequencing with a BrilliantDye Terminator cycle sequencing kit (NimaGen) and an ABI PRISM 3130xl Genetic Analyzer (Applied Biosystems).

The pEGFP-attB vector with the priEE fragment was prepared at a high concentration using Plasmid Midi Kit (Qiagen) and transported to WellGenetics (Taiwan) for injection into two attP strains (*y*^*1*^ *w*^*1118*^; PBac{*y*^*+*^-attP-3B}VK00033 and *y*^*1*^ *w*^*1118*^; PBac{*y*^*+*^-attP-3B}VK00037).

### Deletion strains generated by CRISPR-Cas9

The deletion strains were established by crossing gRNA expression strains with Cas-0002-iso or Cas-0002-iso_*e*::*mCherry* (the mating scheme shown in Fig S2). Deletions (Dels) occur in the germline cells of G1. Twelve G1 males were crossed one by one with several TBX-0010 virgin females. Eight G2 males sampled from the progenies of each G1 male were subjected to PCR screening. DNA samples extracted from the mid-legs of G2 males were amplified using the primers e_-5029F (5’-CGTGTGCCTGATCGCTAGA-3’) and e_-3391R (5’-ACTCGTGCCTTACTTAATCTGAA-3’), which were designed to amplify the target region. The G2 individuals were screened by subjecting the amplicons to electrophoresis using a 1% agarose gel. G2 individuals with deletions were crossed again with TBX-0010. Then, G3 (*y*^*2*^, *cho*^*2*^, *v*^*1*^; +; Del/*TM6c, Sb, Tb*) individuals were crossed with each other to establish G4 homozygous strains (*y*^*2*^, *cho*^*2*^, *v*^*1*^; +; Del). The deletions were confirmed using Sanger sequencing with a BrilliantDye Terminator cycle sequencing kit (NimaGen) and an ABI PRISM 3130xl Genetic Analyzer (Applied Biosystems). The males from the homozygous deletion strains (G4) were crossed twice with the double balancer *w*^*1118*^; *wg*^*Sp-1*^*/ CyO*; *Pr*^*1*^, *Dr*^*1*^*/ TM3, Sb*^*1*^, *Ser*^*1*^ (DGRC#109551) to replace the *y*^*2*^, *cho*^*2*^, *v*^*1*^ X chromosome. Finally, the G7 homozygous deletion strains (*w*^*1118*^; +; Del) were established (Fig S2). Control strains (*w*^*1118*^; +; + and *w*^*1118*^; +; *e*::*mCherry*) were established with the same crosses using TBX-0002 instead of the gRNA expression strain (Fig S2).

### Quantification of pigmentation intensity

At 5–7 days after eclosion, females were placed in 10% glycerol in ethanol at 4°C for 1 h. Next, the flies were rotated in 10% glycerol in phosphate-buffered saline (PBS) at room temperature for 1 h after removing the head, legs, and wings. The images of the dorsal body of flies soaked in 10% glycerol in PBS were captured using a digital camera (DP73, Olympus) connected to a stereoscopic microscope (SZX16, Olympus). The same parameters (exposure time, zoom width and illumination) and reference grayscale (brightness = 128; ColorChecker, X-rite) were applied for capturing all images. White balance was corrected using the white scale (Brightness = 255; ColorChecker, X-rite) with cellSens Standard 1.6 software (Olympus). Pigmentation intensity was measured in manually selected areas of the thorax and abdomen (Figs 2 B and C) from RGB images of flies using ImageJ 1.53a [57]. The mode grayscale brightness values from thorax and abdomen were corrected using the reference grayscale of the background area at the bottom left corner of each image. The percent of darkness was calculated as follows:

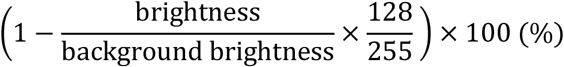

The raw measurement data are in Table S1. The data were analyzed using one-way ANOVA, followed by Tukey HSD post-hoc test. Statistical analyses were performed using R version 4.0.3 [58].

### Confocal microscopy

The adult flies were dissected 4 h after eclosion and the abdomen, wings, front legs, and halteres were collected in PBS. The dorsal abdominal cuticle and epidermis were separated from the rest of the abdomen. The fat body, internal organs, and genitalia were gently removed. The head of adult females collected at 4–4.5 h after the light was turned on was dissected in PBS and the intact brain was obtained. Each brain sample was fixed in 4% paraformaldehyde for 1 h and washed with PBS for 1 h after fixation.

Each specimen was mounted with VECTASHIELD Mounting Medium with DAPI (Vector Laboratories) and imaged under a C2 plus confocal microscope (Nikon). Max intensity images were composited from the XY overlapping images (abdomen: 12 images, wing: 10 images) with 1 μm wide Z-stacks using the NIS Elements AR 4.50.00 software. The following laser wavelengths were applied for obtaining images: 488 nm activation wavelength and 509 nm imaging wavelength for EGFP imaging; 561 nm activation wavelength and 620 nm imaging wavelength for mCherry. The identical parameters of C2 plus settings (HV, offset, laser power, pinhole size, scan size, scan speed, scan direction, and zoom) were applied for imaging the same fluorescence in the same tissue (mCherry or EGFP). No further corrections were applied.

## Supporting Information

**Table S1.** Raw pigmentation scores (10 females per strain) of the strains used in this study.

| strain | Individual<br>number | Thorax |  |  | Abdomen |  |  |
| --- | --- | --- | --- | --- | --- | --- | --- |
|  |  | mode | BG | %darkness<sup>a</sup> | mode | BG | %darkness |
| *w<sup>1118</sup>*; + (control) | 1 | 85 | 161 | 73.5 | 108 | 162 | 66.5 |
|  | 2 | 89 | 162 | 72.4 | 109 | 162 | 66.2 |
|  | 3 | 79 | 163 | 75.7 | 114 | 163 | 64.9 |
|  | 4 | 83 | 164 | 74.6 | 110 | 162 | 65.9 |
|  | 5 | 80 | 162 | 75.2 | 108 | 162 | 66.5 |
|  | 6 | 78 | 161 | 75.7 | 108 | 163 | 66.7 |
|  | 7 | 76 | 162 | 76.5 | 116 | 162 | 64.1 |
|  | 8 | 81 | 162 | 74.9 | 120 | 163 | 63.0 |
|  | 9 | 79 | 162 | 75.5 | 105 | 161 | 67.3 |
|  | 10 | 76 | 161 | 76.3 | 106 | 162 | 67.2 |
| *w<sup>1118</sup>*; *e<sup>Δ1088priEE</sup>* | 1 | 66 | 160 | 79.3 | 119 | 160 | 62.7 |
|  | 2 | 63 | 161 | 80.4 | 120 | 159 | 62.1 |
|  | 3 | 67 | 159 | 78.8 | 131 | 160 | 58.9 |
|  | 4 | 66 | 160 | 79.3 | 133 | 160 | 58.3 |
|  | 5 | 60 | 161 | 81.3 | 121 | 161 | 62.3 |
|  | 6 | 67 | 160 | 79.0 | 124 | 163 | 61.8 |
|  | 7 | 71 | 161 | 77.9 | 133 | 162 | 58.8 |
|  | 8 | 64 | 161 | 80.0 | 111 | 163 | 65.8 |
|  | 9 | 67 | 160 | 79.0 | 122 | 162 | 62.2 |
|  | 10 | 63 | 161 | 80.4 | 119 | 161 | 62.9 |
| *w<sup>1118</sup>*; *e<sup>Δ1089priEE</sup>* | 1 | 66 | 159 | 79.2 | 117 | 160 | 63.3 |
|  | 2 | 68 | 160 | 78.7 | 112 | 159 | 64.6 |
|  | 3 | 66 | 160 | 79.3 | 108 | 163 | 66.7 |
|  | 4 | 64 | 161 | 80.0 | 107 | 160 | 66.4 |
|  | 5 | 69 | 160 | 78.4 | 117 | 160 | 63.3 |
|  | 6 | 68 | 160 | 78.7 | 109 | 160 | 65.8 |
|  | 7 | 64 | 160 | 79.9 | 119 | 161 | 62.9 |
|  | 8 | 71 | 161 | 77.9 | 116 | 161 | 63.8 |
|  | 9 | 65 | 161 | 79.7 | 133 | 160 | 58.3 |
|  | 10 | 68 | 161 | 78.8 | 117 | 163 | 64.0 |
| *e<sup>1</sup>* | 1 | 25 | 159 | 92.1 | 88 | 159 | 72.2 |
|  | 2 | 24 | 161 | 92.5 | 86 | 159 | 72.8 |
|  | 3 | 23 | 160 | 92.8 | 76 | 158 | 75.9 |
|  | 4 | 22 | 161 | 93.1 | 70 | 159 | 77.9 |
|  | 5 | 23 | 160 | 92.8 | 69 | 159 | 78.2 |
|  | 6 | 23 | 161 | 92.8 | 70 | 159 | 77.9 |
|  | 7 | 25 | 158 | 92.1 | 73 | 159 | 77.0 |
|  | 8 | 23 | 159 | 92.7 | 74 | 160 | 76.8 |
|  | 9 | 21 | 160 | 93.4 | 76 | 161 | 76.3 |
|  | 10 | 24 | 160 | 92.5 | 76 | 159 | 76.0 |

mode: grayscale mode of quantified area; BG: gray background area
<sup>a</sup> % darkness was calculated from mode and BG.

**Fig S1.**
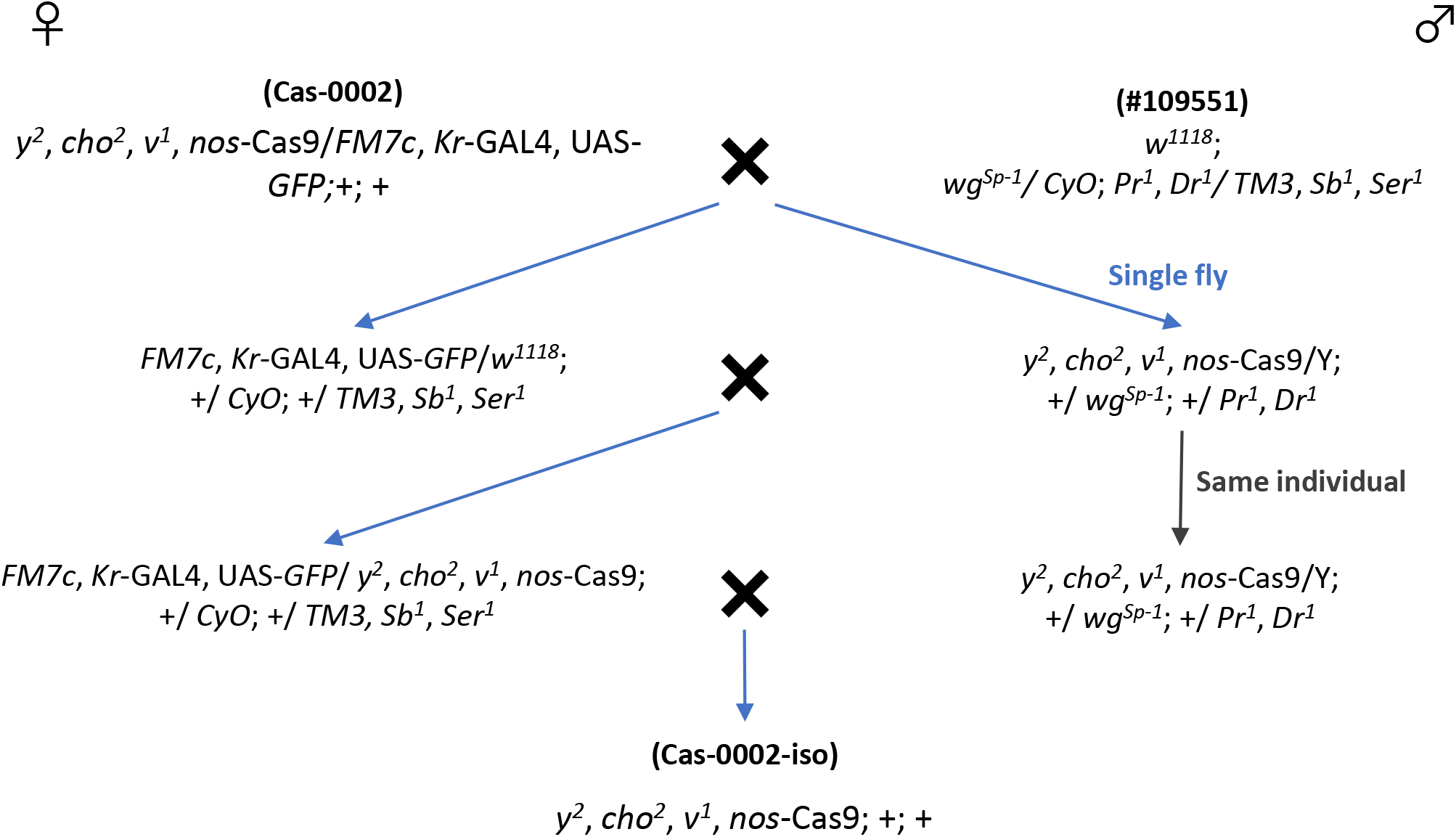
Scheme for extracting an isogenized chromosome. To control for the genetic background of the genome-edited flies, each chromosome was originated from a single chromosome. The isogenized strain was named Cas-0002-iso.

**Fig S2.**
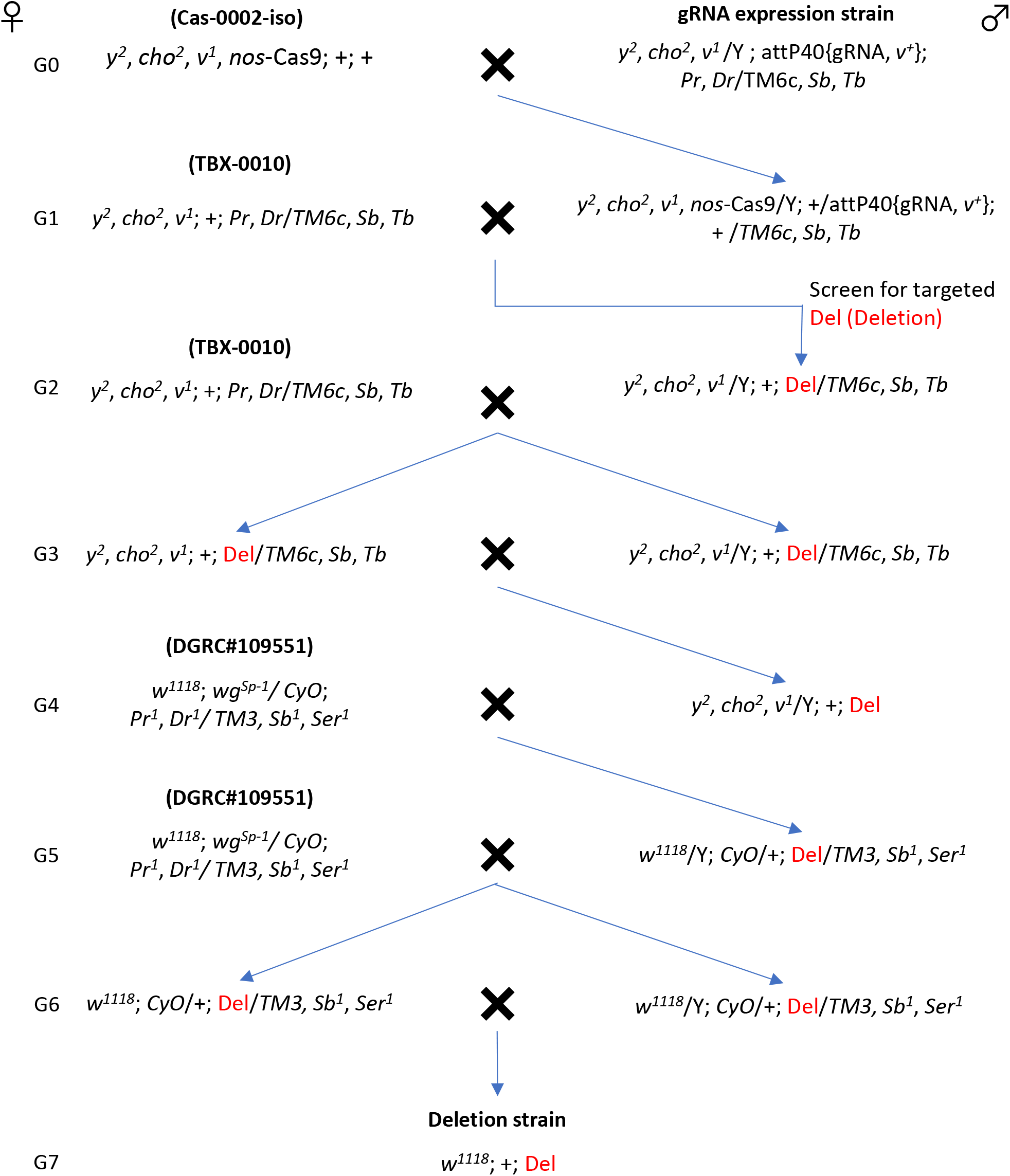
Scheme for generating primary enhancer element (priEE) deletion strains without *y*^*2*^. CRISPR-Cas9-based genome editing was performed by crossing the Cas-0002-iso with the guide RNA (gRNA) expression strain (G0). The progenies from the G1 cross were screened for the presence of deletions. Homologous deletions were achieved by the crosses in G3 and G4. *y*^*2*^ was removed by the crosses in G4 to G7 because it interferes with *ebony* in the pigment biosynthesis pathway. Deletion strains for mCherry fluorescence observation were established using the same scheme, except Cas-0002-iso_*e*::*mCherry* was used instead of Cas-0002-iso (G0). Control strains (*w*^*1118*^; +; + and *w*^*1118*^; +; *e*::*mCherry*) were established with the same crosses using TBX-0002 instead of the gRNA expression strain and using + instead of the Del genotype for the G2 cross.

**Fig S3.**
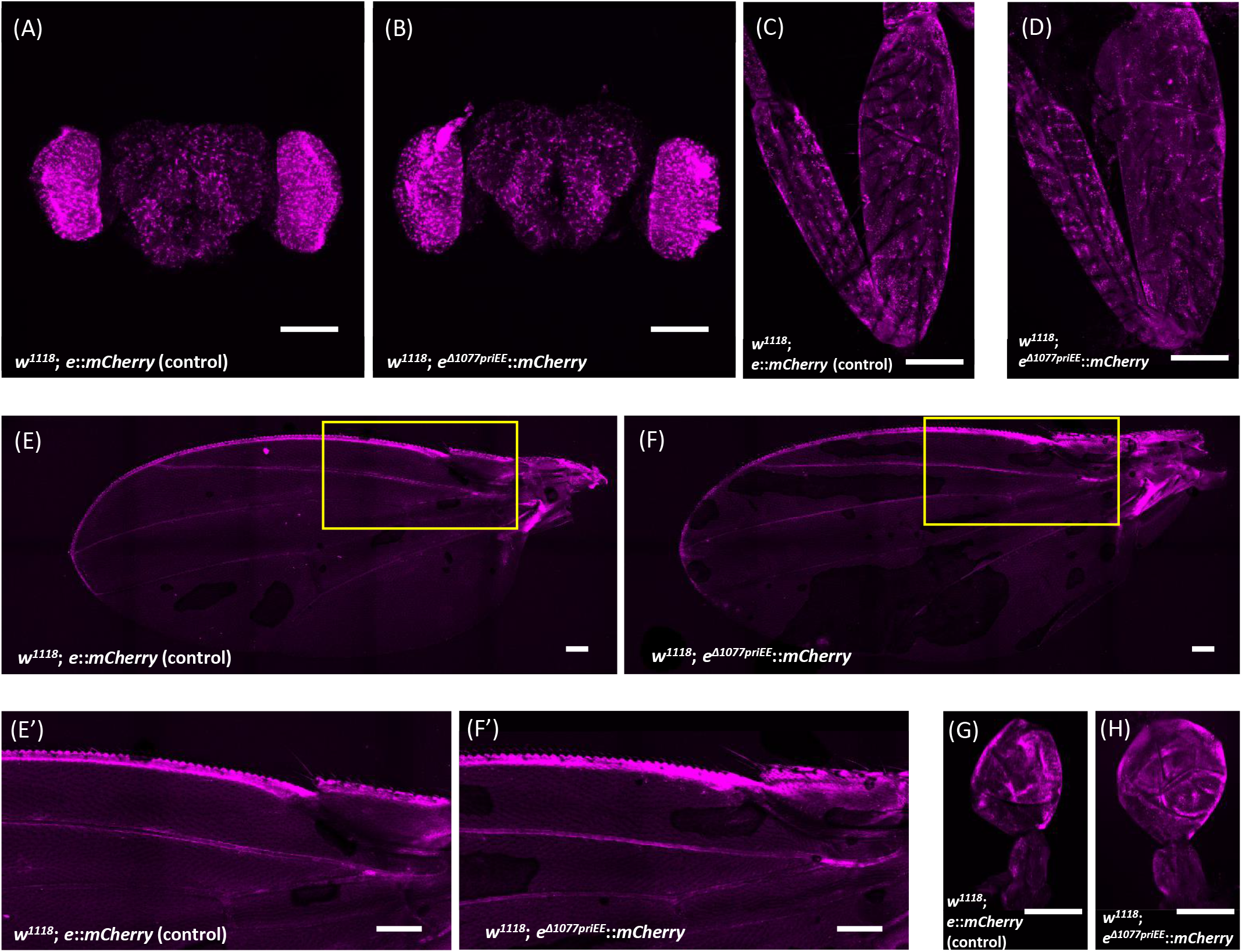
*ebony* expression in tissues other than developing epidermis. Confocal images of brain (A and B), front leg (C and D), wing (E, E’, F and F’), and haltere (G and H). Each tissue was dissected from the control, *w*^*1118*^; *e*::*mCherry* (A, C, E, E’ and G), and the priEE-deleted strain *w*^*1118*^; *e*^*Δ1077priEE*^::*mCherry* (B, D, F, F’ and H). (E’) and (F’) are magnified views of the yellow square in (E) and (F), respectively. The scale bars indicate 100 μm.

**Fig S4.**
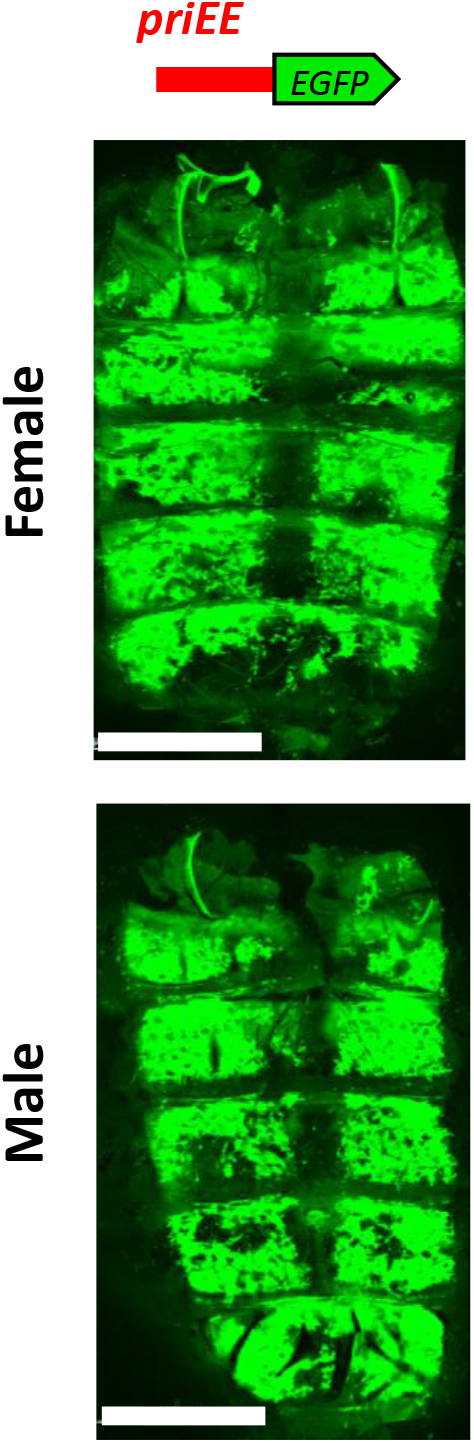
Reporter assay using the enhanced GFP-tagged primary enhancer element (*priEE-EGFP*) construct in the developing abdominal epidermis. Confocal fluorescence images of the developing abdominal epidermis of *priEE*-*EGFP* transformed to VK00037. Images of females and males are shown in the upper and lower panels, respectively. Scale bars indicate 500 μm.

**Fig S5.**
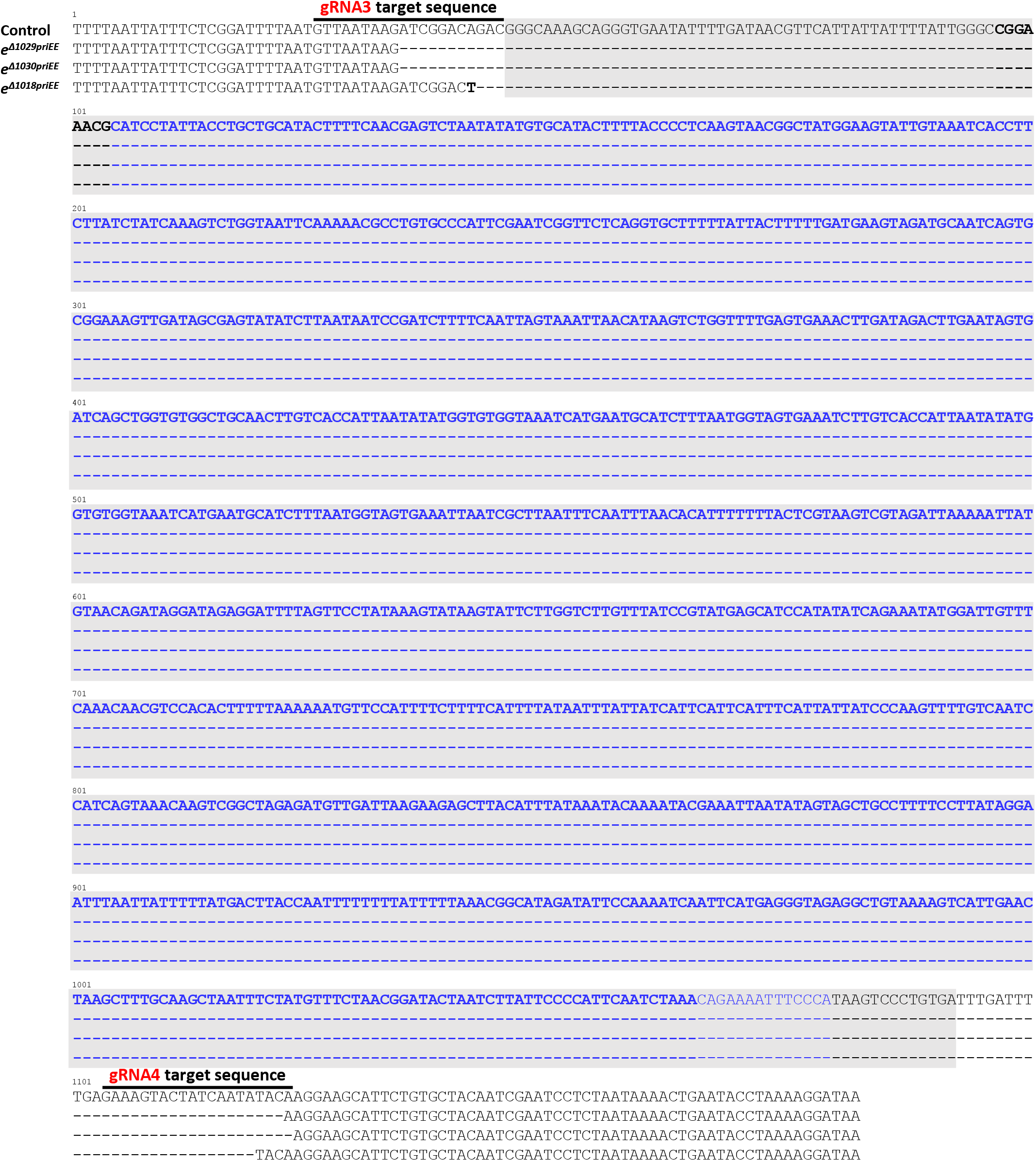
Alignment of sequences around the primary enhancer element (priEE) in the control and priEE knockout strains. The guide RNA (gRNA) target sequences are indicated above the sequence. Bases in bold indicate the 947-bp sequence of *e*_ECR0.9 from Takahashi and Takano-Shimizu [29] (969-bp sequence in the control strain (Cas-0002-iso-derived *w*^*1118*^; + strain)). Bases in blue indicate the 961-bp sequence of *e*_core_*cis* sequence from Miyagi *et al*.[28] (975 bp in the control strain (Cas-0002-iso-derived *w*^*1118*^; + strain)). Shaded bases indicate the sequence fragment (1,047 bp) used for the reporter assay in this study.

## Acknowledgment

We would like to thank Nikolas Gompel for *pJet-yellow_F4mut-mCherry* vector, WellGenetics for embryonic injection, and NIG-Fly and Kyoto Stock Center for fly stocks. We would like to thank Koichiro Tamura and the Evolutionary Genetics Laboratory members at TMU for critical comments and discussions. The work was partly supported by the Sasakawa Scientific Research Grant (No. 2019-4038) from The Japan Science Society and by JSPS KAKENHI (Grant No. JP19H03276) awarded to N.A. and A.T., respectively.

## Author contributions

Conceived and designed the experiments: NA KMT AT.

Performed the experiments: NA SS.

Analyzed the data: NA SS.

Contributed reagents/materials/analysis tools: KMT TS AT.

Wrote the paper: NA AT.

